# Seasonal plasticity and energy efficiency in migratory buntings is an assemblage of alterations in metabolic and gut microbe adaptations

**DOI:** 10.1101/2023.02.17.529037

**Authors:** Neelu Jain Gupta, Samya Das, Mrinal Das, Rakesh Arya, Anit Kumar, Ranjan Kumar Nanda

## Abstract

Circulatory system is the source of useful metabolites to enable organismal energy demands of muscle during different life states of birds including migration. In this study, we profiled the serum metabolites and fecal microbiome of redheaded buntings (*Emberiza bruniceps*) exhibiting diurnal non-migratory (*nonM*) pre-migratory (*preM*), migratory (*M*) and post-migratory (*posM*) states, when exposed to long days (14L:10D). Using gas chromatography, out of the identified 124 serum analytes, 38 showed significant variations (Fold change, VIP) between states. Out of these, 11 metabolites (short chain fatty acids, SCFAs-butanoate and hexanoate, lactate, pyruvate, ethylene and acetate oxime, pyridoxal phosphate (PLP), niacin, L-valine and carboximidic acids viz. phosphatidyl ethanolamine (PE) and diethyl carbamate) involved in energy pathways and enhanced immunity showed higher abundance in *M* state. While, upsurge of L-valine suggested energy contribution of glucogenic branched chain amino acids (BCAAs) in *M* state, that of leucine metabolite, was related to higher temperature in *posM* birds. Gut microbiota was analysed using faeces of buntings (n=6 each) during nonM and M states and alteration in bacterial compositions was observed; faeces of nonM birds were enriched in proteobacteria, while those of M were rich in firmicutes. This study reports the migratory state specific changes in the serum metabolome and faecal microbiome of the buntings and highlights the role of short chain fatty acids, SCFAs and branched chain amino acids, BCAAs during hypermetabolic state of migration.

## Introduction

Every year, migratory birds perform magnanimous feat of spring and autumn migrations, exhibiting nonstop night flights which varies from few days to two months. Triggered by changes in natural daylength, regulated by circannual clock, wintering nonmigratory birds (*nonM*) go through premigratory preparation state (*preM*) before undertaking spring migratory (*M*) night-flight. Phenotypically, intense migratory exercise, accrues subsequent weight loss in the post-migratory state (*posM*). Migratory birds adopt strategies to accumulate and switch energy substrates, expedite lipogenesis, and transport fatty acids to muscles to achieve continuum of extreme metabolic transitions from *nonM, preM, M* up to *posM*, while limiting unfavourable metabolic consequences (Mcfarlan, 2009; Guglielmo, 2010). For example, during migration, hyperglycaemic (Szwergold, 2014) northern wheatears’ adaptively replace glucose with lipid substrates to improve energy homeostasis, whereas in captive birds gives metabolic solution to defatting (Frias-Soler et al., 2021). Tiny Ruby-throated hummingbird are reported to avoid glucosuria through higher turnover rates of red blood cells and proteins (Hargrove, 2005). Even the seasonal switching of food type available in nature maximizes carbon and nitrogen content to increase dietary amino acids intake (Gómez, 2018). Although, investigations on migratory syndrome included migratory physiology (Butler, 2016), sleep deprivation (Horton et al., 2019), hormonal regulations (Mishra et al., 2017), fat mobilisation (Guglielmo, 2010) and others, limited studies on the role of metabolic components of these states during migration are reported.

Night migratory songbird, redheaded buntings (*Emberiza bruniceps*), winter visitors to Indian subcontinent from breeding ground (40°N) and live a lean (*nonM)-obese(preM)-* superobese(*M*)-lean(*posM*) annual life. Comparative transcriptomic signatures of buntings (Sharma et al., 2019, 2021a), also suggest differential acetyl-CoA requirement, between the *nonM/M* states, reinforcing importance of metabolites leading to energy efficiency from non-glucogenic sources like lipids and amino acids. Recently, omics analyses engaging tissues specific proteome and metabolite expressions (Banerjee & Chaturvedi, 2016, Gupta et al., 2020; Sharma et al., 2021b) suggested the role of fatty acids (FA) metabolism in migratory buntings. Alteration in expression of candidate genes (Sharma et al., 2021) during heightened activity in brain in mediobasal hypothalamus, MBH, liver and muscle suggest increase in oxidative stress at cellular level (Sharma et al., 2018; Valek et al., 2019). Circulatory biofluids like blood provides a rich metabolic components draining from different organs to complete inter-organ distribution and maintain homeostasis. We aimed to capture the variation in the serum metabolic components during different photoinduced *nonM, preM, M and posM* of spring migration in the annual life history states (LHS) of redheaded bunting (Figure 1A). The important “serum biomarkers” could explain the metabolic phenotypes that exist in different migratory states of the model system i.e. buntings.

**Figure 1.**
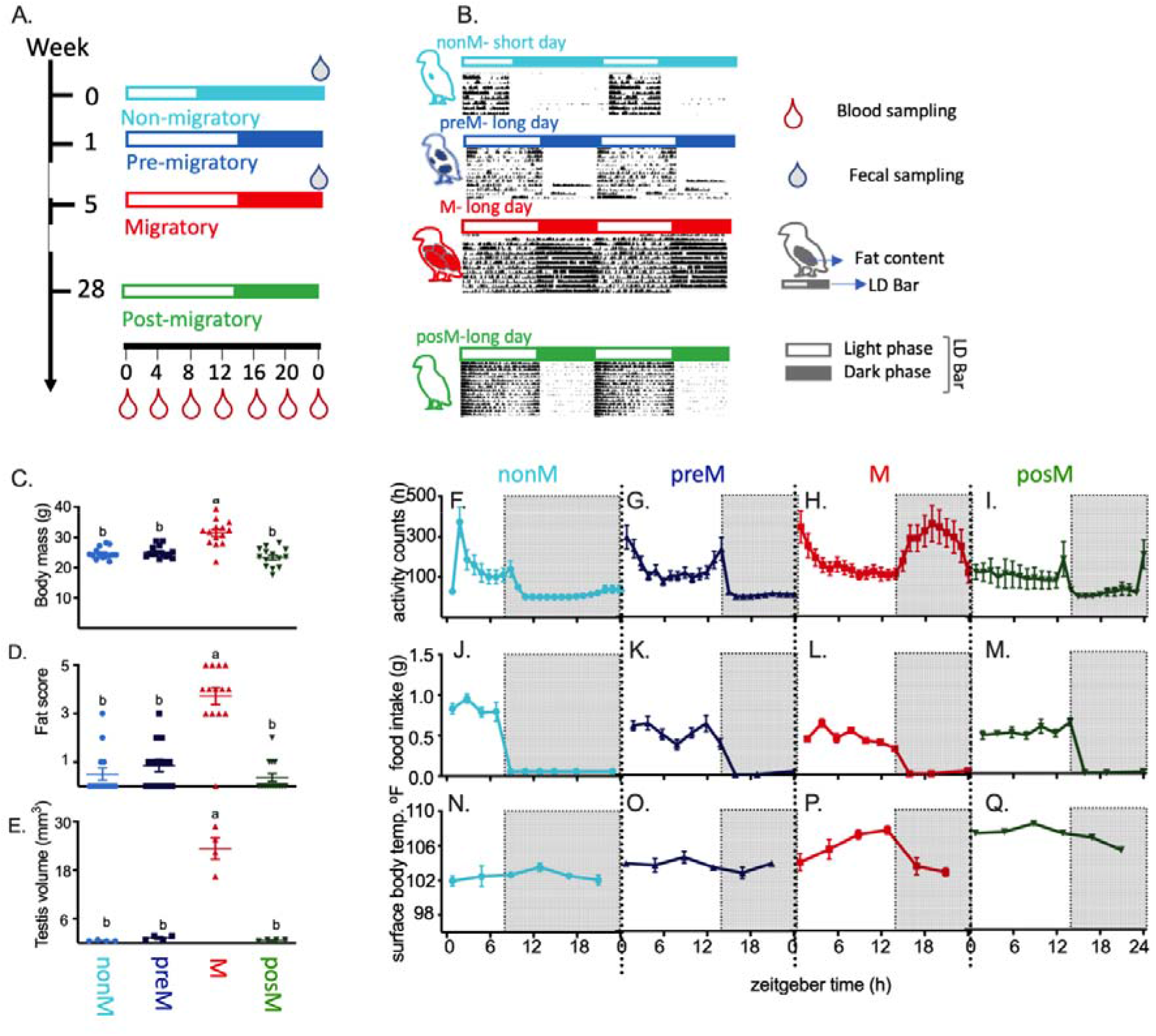
Experimental design adopted in this study to identify serum based metabolites in migratory states in redheaded buntings. **A.** Groups of photosensitive redheaded buntings held under short days (8h light:16h darkness, SD, nonM, cyan blue) were either continued in SD or transferred to LD (14h light:10h darkness), on day 0 and sampled at 1 (preM, blue), 5 (M, red) and 28 (posM, green) week intervals. Blood was drawn at 4-hourly intervals (hollow red drops on time scale) treating lights on as 0 ZT (zeitgeber time), hollow blue drops show time of fecal sample collection, open and closed bars indicate light and dark phases, respectively. **B.** Representative double-plotted activity record (actograms) of birds held under SD and LD as described in 1A. Extent of fat deposition is indicated separately. Changes in body mass in grams (**C**), fat score (**D**) and testis (**E**) volume (mm^3^) during nonM, preM, M and posM life history states. Daily profiles of changes in activity (**F-I**), food consumption (**J-M**), and surface body temperature (**N-Q**), in nonM, preM, M and posM buntings. (nonM-nonmigrating; preM-premigratory; M-migratory and posM-postmigratory birds).

## Results

We employed global serum metabolite analysis to monitor the differences in energy budgeting during different states of migration in buntings. Different states of migration of redheaded buntings (*Emberiza bruniceps*) i.e. *nonM, preM, M and posM* showed differences in their behavior and physiological parameters (Figure 1B–1D). As expected, buntings were day active in *nonM, preM* and *posM* states, while exhibiting temporal behavioural shift resembling *zungunruhe* in wild conspecifics in *M* state (t_26_= 2.317, P < 0.05; Figure 1B, 1H). Significant variation in the behavioural activities during the three non-migrating states i.e. *nonM, preM* and *posM*, with respect to time of day (F_23, 552_= 9.162, P < 0.0001; Figure 1B, 1F–1I) were observed whereas it was absent in the *M* state (F_3,24_= 1.7 68, P > 0.05). A bimodal daily food eating pattern (DFEP) through the light phase in *nonM* (Figure 1J), *preM* (Figure 1K) and *posM* (Figure 1M) states were observed which was missing in *M* state (Figure 1L). During the non-migrating states (*nonM, preM* and *posM*) the total daily food intake was low but high in *M* state. Surface body temperature of *posM* (Figure 1Q) birds was significantly higher (F_15,90_= 2.594, P < 0.05) than *nonM, preM* and *M* (Figure 1N–1P). The extreme transitions in physiological activity may influence in their serum metabolic component and might be useful to understand their phenotypes.

A total of 144 serum samples collected longitudinally at 4-hourly intervals from birds (n=6, Figure 1A) of different states i.e., *nonM* (0 week), *preM* (1 week), *M* (5 weeks) and *posM* (28 weeks) from the transfer to photo-inducing daylengths. Global metabolite analysis of these serum samples was run in batches of 20 using GC-MS and out of the identified molecular features (n=1233), 124 selected for multivariate analysis using MetaboAnalyst 4.0. A set of 38 deregulated important molecules (variable projection parameter (VIP) score>0.1) were identified between *M* and other states (*nonM, preM, posM*). Principal component analysis (PCA) of the serum analytes showed LHS specific clusters (Figure 2A, 2B).

**Figure 2.**
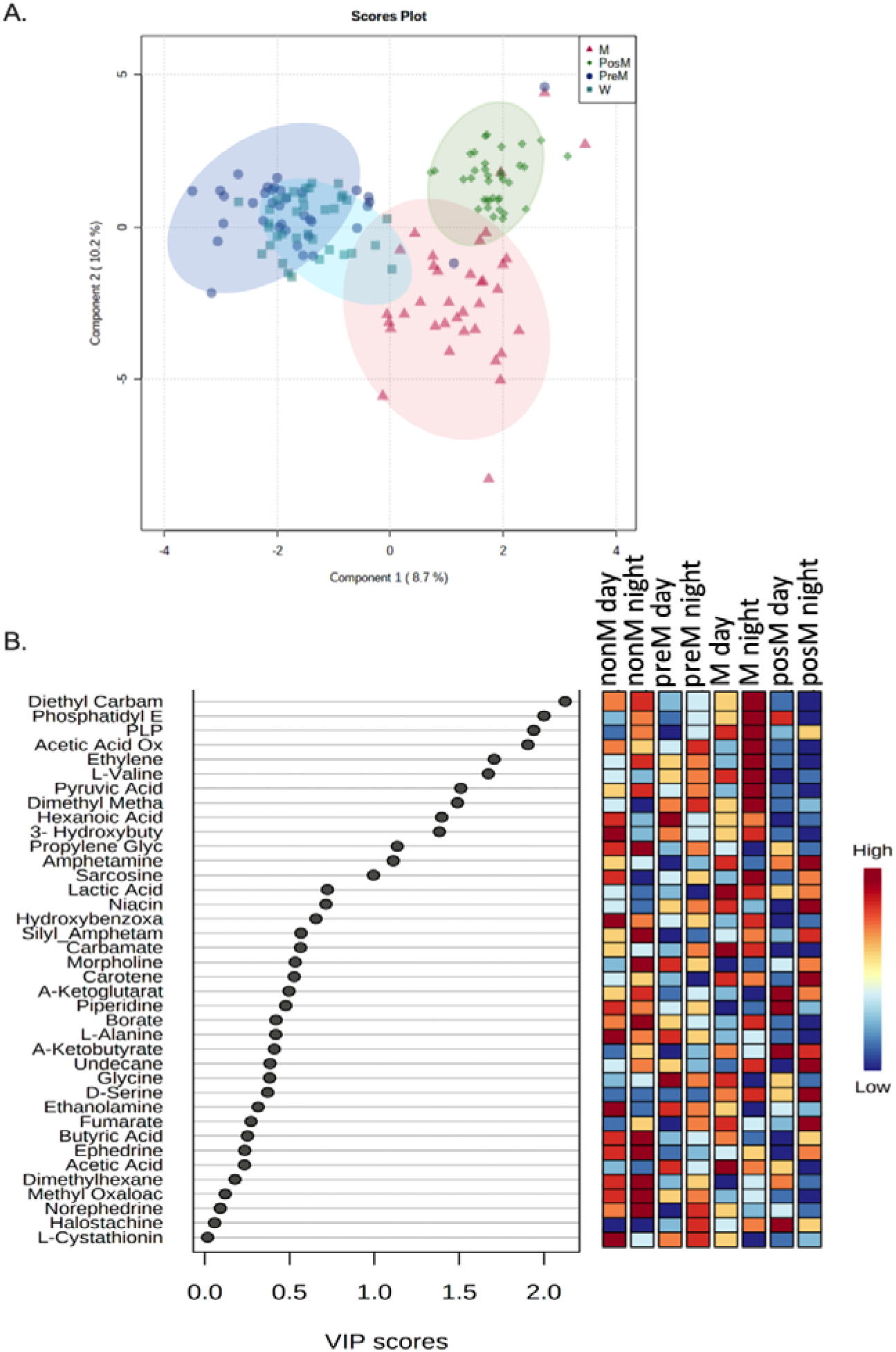
Serum metabolite analysis of birds at different states of migration. **A**. Principal Component Analysis (PCA) of serum analytes of nonM (nonmigrating, cyan blue), preM (premigratory, blue), M (migratory, red) and posM (postmigratory, green) showed state specific clustering. **B.** Partial Least Square Discriminant (PLS-DA) model built using serum analytes from day and night serum of redheaded buntings photoinduced to nonM, preM, M and posM life-history states. Analytes with variable importance in projection (VIP) score >0.1 of PLS-DA are shown.

Diurnal trend of the identified state specific important serum metabolites in buntings are presented (Figure 3 and Supplementary Figure S1). Serum lactate levels exhibited inter-LHS variations (F_3,24_= 7.73, P < 0.0005) with highest serum levels in the preM and M. Serum pyruvate levels showed inter-LHS variations (F_3,20_= 5.293, P < 0.0075) and mean *nonM* (t_10_= 3.019, P < 0.05), *preM* (t_10_= 3.323, P < 0.01) and M (t_10_= 3.235, P < 0.01) levels were significantly higher than *posM*. Serum acetate levels in buntings exhibited inter-LHS variations (F_3,20_= 3.659, P < 0.05), were higher in preM (t_10_= 2.9, P < 0.05) and M (t_10_= 4.24, P < 0.005) than *posM*. TCA cycle intermediates like alpha keto-glutaric acid (F_3, 19_= 0.65) and fumarate (F_3, 20_= 0.35) showed similar levels in inter-LHS. Serum SCFAs like hexanoate (F_3,20_= 8.709, P < 0.05) and butanoate (F_3,20_= 6.44, P < 0.05) varied during inter LHS, exhibiting higher levels at night of *M* state compared to others (*nonM, preM* and *posM*). Methyl oxalacetate, glutathione synthesis transamination intermediate, was significantly lowered during night of M and posM states (F_3,20_= 13.61, P < 0.0001).

**Figure 3.**
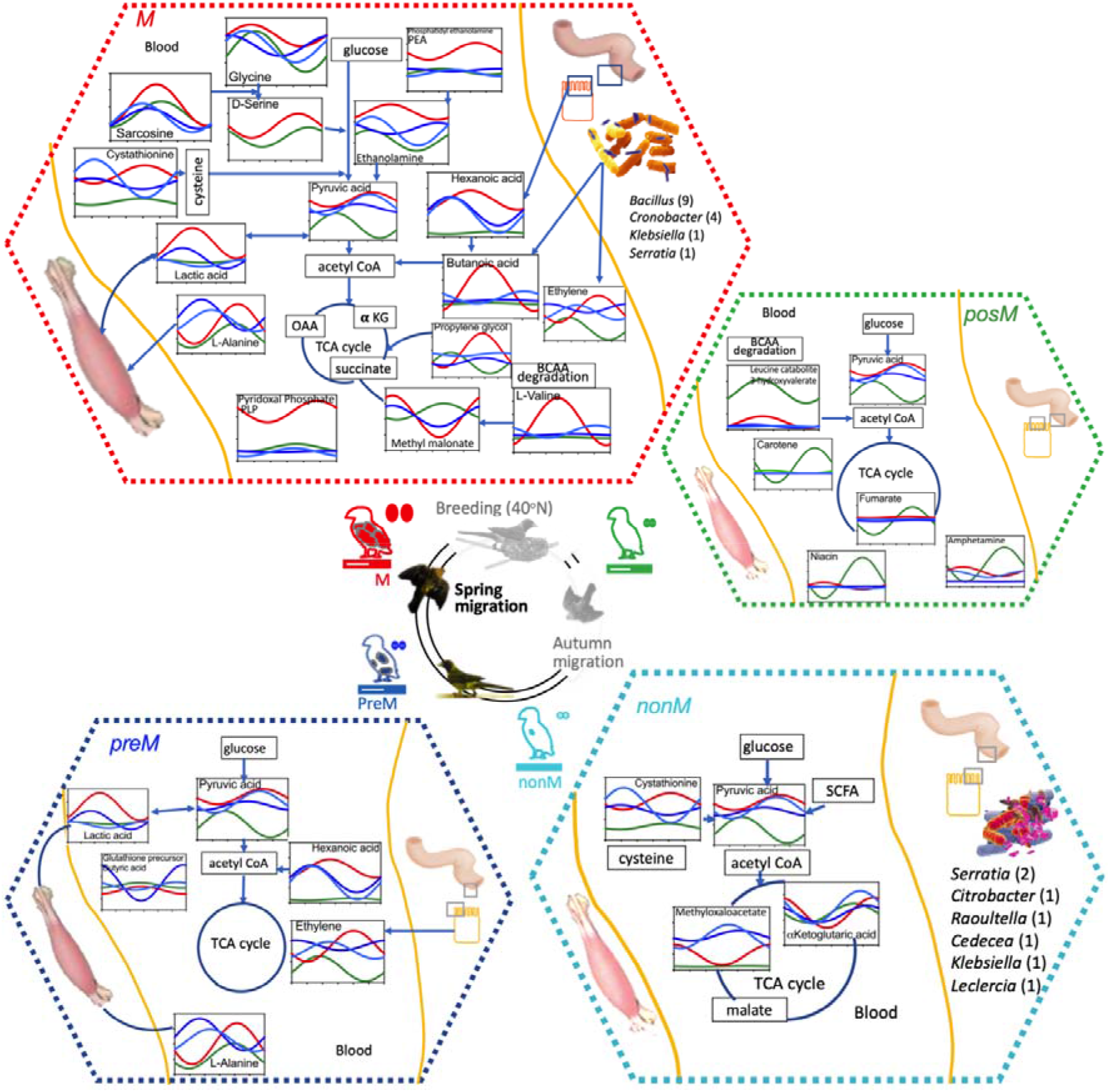
The central circle shows annual life history states of redheaded bunting, highlighting spring migration in colour. All insert boxes show Cosinor curves to depict relative profiles of serum metabolite concentrations during nonM (cyan blue, non-migratory), preM (blue, premigratory), M (red, migratory) and posM (green, postmigratory) states. Circulating metabolites are shown in broken hexagon boxes on the basis of higher abundance during specific LHS and gut microbiota composition during non-M and M states are shown separately.

Pyridoxal Phosphate (PLP) coenzyme, active form of vitamin B6, was significantly (F_3,20_= 18.86, P < 0.0001) higher in *M* state. Daily (F_5,108_= 34.76, P < 0.0001; F_5,118_= 7.903, P < 0.0001) and inter-LHS (F_3,108_= 65.6, P < 0.0001; F_3,118_= 14.05, P < 0.0001) variations were highly significant in two vitamins, carotene and niacin (Figure 3).

Phosphatidyl ethanolamine (PE), phospholipid plasmogen/ chaperone in mitochondrial pathway/ activity exhibited significant seasonal variations (F_3,20_= 16.17, P < 0.0001). PE precursor carboximidic acid and ethanolamine was found to be higher in serum of birds of *M* state (F_3,20_= 3.894, P < 0.05).

A set of amino acids (valine, serine, alanine, glycine) and amino acid degradation intermediates (sarcosine, propylene glycol, 3-hydroxy valerate, cystathionine, α-hydroxybutyrate) were detected in the serum of these study groups (Supplementary Figure S2). Higher levels of serum valine (F_3,20_= 3.149, P < 0.05) and serine (t_10_= 1.83, P < 0.0479) were observed in *M* group. Serum alanine levels were higher (t_10_= 2.635, P < 0.05) during *preM* compared to *posM* but remained similar in non*M* and *M* group. Serum glycine level was found to be similar between states and did not exhibit inter-LHS variations. However sarcosine which is an intermediate from choline to glycine conversion showed an inter-LHS (F_5,120_= 7.6, P < 0.0001) and within group (F_3,120_= 4.6, P < 0.005) daily variations. An intermediate of isoleucine i.e. propylene glycol (F_3,20_= 0.2578, P= 0.8549), found to be similar between states with minor daily variation in LHS. Ketogenic degradation intermediate of leucine i.e. 3-hydroxy valerate, showed 16 fold higher abundance in *posM* (F_3,20_= 7.92, P < 0.001). Serine-cysteine pathway intermediate cystathionine showed a minima in *posM* (F_3,20_= 6.214, P < 0.005). α-hydroxybutyrate (herein, butyric acid) is implicated in glutathionylation (L-γ-glutamyl-L-cysteinyl-glycine) of cysteine. This trans-sulfuration/ transamination intermediate, showed 4-fold higher abundance during ZT12 at *posM*, thus exhibiting significant (F_5,69_= 2.74, P < 0.05) daily variations (Supplementary Figure S1). Ornithine synthesis intermediates, carbamic acid and diethyl carbamate, implicated in nitrate reduction, exhibited inter-LHS variations (F_2,15_= 8.089, P < 0.005 and F_3,20_= 9.595, P < 0.0005, Supplementary Figure S1, S2).

### Gut microbiota differs among non-migratory and migratory LHS

Bacteria identified in bunting faecal samples (Figure 3 and Supplementary Figure S3), comprised two families of phylum Proteobacteria (gram negative; Enterobacteriaceae and Yersiniaceae) and one family of phylum Fermicutes (gram positive; Bacillaceae). Bacterial species common to both LHS included *Serratia liquefaciens, Cedecea lapagei*, and *Klebsiella aerogenes*. Bacteria identified from non-migratory bird faeces included *Raoultella ornithinolytica, Serratia marcescens, Leclercia adecarboxylata and Serratia nematodiphila*. Migratory bird faeces contained 9 non-pathogenic species of *Bacillus* viz. *xiamenensis, aerophilus, stratosphericus, zhangzhouensis, aerius, australiameris, altitudinis, safensis, pumilus* and 4 species of *Cronobacter* viz. *turicensis, muytjensii, dublinensis* and *sakazakii*.

## Discussion

The study is first report on daily orchestration of serum metabolites during different photoperiod-induced (a simulation of spring migration) LHSs in a migratory songbird.

### Inter-LHS and diurnal variations in serum metabolites of redheaded buntings

During the night cycle of *M* state, experimental birds exhibited wing whirring activity and a state of energy metabolism, similar to migrating wild conspecifics. 38 of the 124 serum analytes identified using GCMS, showed significant variations. During *M* state, 11 metabolites showed higher abundance, which included energy metabolites like SCFAs viz. butanoate and hexanoic acid; energy pathway intermediates viz. lactate, pyruvate, ethylene and acetate oxime; vitamins viz. PLP and niacin; amino acid, L-valine; carboximidic acids viz. PE and diethyl carbamate.

### Inter-LHS differences in energy metabolites also draw attention to enhanced immunity

During endurance flight, bunting needs to quickly transport energy-dense metabolic fuels like liver fat to muscles (Butler 2016), substantiated by high FAT/CD36 and FABP (fatty acid binding proteins) genes expression in muscle (Guglielmo, 2010). Higher SCFA levels in serum of *M* state of buntings’ support SCFAs as breakdown intermediates, transported through blood during high energy demands. In poultry, SCFAs inhibit pathogenic invasion and colonization by lowering the intestinal pH, besides promoting the differentiation of T cells into T regulatory cells and expansion by binding to receptors such as Toll-like receptors (TLR) and G protein-coupled receptors (GPCRs) on immune cells (Liu et al., 2021). SCFA abundance appears to be associated with microbial degradation (Kim et al., 2019) in *M* buntings. Notably, Skeen et al., 2021 have also reported alteration in microflora of several migratory birds’ faeces. The possibility of increased *SCFA* abundance in M buntings being related to increased microflora, cannot be ruled out. Contrary to expectation, we found state specific inter-LHS variation in energy metabolism and Krebs’s cycle intermediates (pyruvate and acetic acid) during *M* state. Similar findings in the hypothalamus and liver of buntings showed seasonal alternations in gene expression of pdc and pdk that mediates pyruvate to acetate conversion (Trivedi et al., 2015).

### Inter-LHS differences in metabolites implicated in reactive species and immunity

Besides, SCFAs, phosphatidyl ethanolamine (PE) and ethanolamine (EA) also exhibited altered abundance in *M* state of the buntings. Firmicutes and some Bacteroidetes were abundant in gut microbiota of migratory shorebirds (Zhang, 2021). PE localised in mitochondria (Sam et al., 2021) also mediates PPARα associated stimulation of FA oxidation, also reported in spring migrating gray catbirds (Corder et al., 2016). EA was reported to have anti-inflammatory and cannabinomimetic properties (Gupta et al., 2020). Higher levels of ketone metabolites like β-hydroxybutyrate, capable of downregulating inflammasome of the immune system, in the *M* state corroborated earlier findings (Horton et al., 2019). During *M* LHS, higher serum lactate levels suggested hyper-metabolic activity driven by oxidative stress. The hyperlacticaemia also drew attention to gut inhabiting Firmicutes during *M*, however it needs detailed investigation for confirmation in future studies. Higher serum PLP supports higher antioxidant ability.

### Inter-LHS differences in amino acids metabolism

Elevated levels of L-valine and isoleucine intermediate i.e. propylene glycol in *M* state suggest an upsurged energy contribution of glucogenic BCAAs. Intermediates of the Leucine metabolism i.e. 3-hydroxy valerate was higher in *posM* state. Ketogenic BCAA leucine (in adipose tissue and muscle), is either degraded via cellular regulatory molecule acetate or incorporated into body protein synthesis under stress conditions like fasting (Rosenthal et al., 1974). Herein, we hypothesize that during *posM* stressed condition, considering lower acetate levels, leucine metabolism is redirected to protein synthesis (Kim et al., 1996). Higher leucine levels reported to be associated with higher body temperature to provide/ thermostability in broiler chicks (Han et al., 2019). Although metabolic fate of L-leucine in *posM* in blood is not known, higher body temperature of *posM* buntings might relate to higher leucine intermediate levels. Higher diethyl carbamate serum influx (also see Supplementary Figure S1) derived from cystathionine-glutathione metabolism, alongside higher methyl oxaloacetate levels (transamination intermediate from glutathione biosynthesis and precursor to asparto-arginine) suggested anaplerotic amino acid energy contribution, during *M* state *However, protein degradation is also subject to fermentation efficiency of gut microbiota* (Gilbert et al., 2018).

Alanine is produced by reductive amination of pyruvate and reversibly formed by aminotransferase action on glutamate, has key role in glucose homeostasis (Cooper & Jeitner, 2016), this explains high levels of serum alanine during increasing energy state of *preM*. Carbamates, substrate for glucuronidation was also high during *M* state, indicating semi-captive bird’s escalated symptomatology to dietary pesticides (Walker, 2003).

### Gut microbiota differentiated obese exercising migrants from lean non-migrants

The species of *Serratia, Cedecea* and *Klebsiella* found in faeces of buntings, during both, migratory and non-migratory states (Figure 3, Supplementary Figure S3) were implicated in metabolite breakdown, such as, *Serratia liquefaciens* (Joshi & Kozlowski, 1987) metabolizes lysine and ornithine for carbon source and energy, *Cedecea lapagei* (Thompson & Sharkady, 2020) was involved in lipid emulsification and Klebsiella’ *aerogenes* (Wesevich et al., 2020) metabolizes citric acid to produce pyruvic acid. Bacteria identified from *nonM* bird faeces i.e. *Raoultella ornithinolytica* (Hajjar et al., 2020)*, Serratia marcescens* (Greenberg, 1978)*, Leclercia adecarboxylata* (Keyes et al., 2020) *and Serratia nematodiphila* (Zhang et al., 2009) were largely facultative anaerobic. *Raoultella ornithinolytica* is an important food regulator (Yoshimatsu et al., 2002), reduces birds’ appetite and might play role in lower food consumption during *nonM* (Jain & Kumar, 1995). *Serratia marcescens* reported to mediate oxidative lactic acid production and citrate degradation in gut, whereas *Serratia nematodiphila* utilizes lactate, l-ornithine and proline. (Zhang et al., 2009) *Leclercia adecarboxylata* (Keyes et al., 2020) utilizes malonate for carbon source and corroborates with our observation of lower malonate levels in *nonM*. Migratory bird faeces included 9 non-pathogenic species of Firmicutes i.e. *Bacillus spp*., mainly implicated in lipase activity and fat digestion. Most *Bacillus spp*. identified such as *B. xiamensis* (Lai et al., 2014), *B. aerophilus* (Branquinho et al., 2015), *B. stratosphericus, B. aerius* (Shivaji et al., 2006), *B. australimaris, B. zhangzhouensis, B. pumilus* (Liu et al., 2016)*, B. altitudinis* (Mao et al., 2013) and *B. safensis* (Singh et al., 2003) found in migratory faeces embroil probiotic activity, which might be directly related to increased metabolic efficiency during migration. Although flying animals are extremely conservative in harbouring enormous microbes (Song et al., 2020). Our study, for the first time suggests that gut microbiota is important in buntings’ metabolic efficiency during migration. This suggestion corroborates with findings on non-obese and obese rats, which, when exposed to controlled exercise, exhibited differences in type and abundance of gut microbiota, alongside alteration in serum lactate levels (Petriz et al., 2014). Gut microbiota might also be important in reactive species scavenging. *Cronobacter turicensis*, known to reduce reactive species (Stephan et al., 2011), produces enzyme superoxide dismutases (SOD) that reduces toxic oxygen radicals. *Cronobacter muytjensii* (Iversen et al., 2008) mediates reductive deamination, producing pyruvic acid and energy in presence of pyridoxal phosphate.

Overall, we report for the first time, the differences in metabolic phenotypes during migratory-nonmigratory states, in redheaded buntings. Buntings exhibit adaptation in pathway intermediates, related to energy metabolism and gut microbiota. Our study highlights that obese spring migrants adaptively differ from non-obese nonmigrating state buntings in their serum energy metabolites levels and gut microbiota. This opens avenues for future investigation to correlate contribution of higher serum lactate levels in hypermetabolic state of migration and possible role of gut microbiota.

## MATERIALS AND METHODS

Experiment was performed on male redheaded buntings, *Emberiza bruniceps*, following the Institutional Animal Ethics Committee (IAEC) guidelines of Chaudhary Charan Singh University, Meerut. Birds obtained from overwintering flocks in February were acclimatized in aviary for 3 weeks. Food (*Setaria* seeds with boiled egg) and water were provided *ad libitum*.

Buntings (n= 48) were housed indoors in well-aerated photoperiodic chambers (size: 1 m × 1 m × 1 m) under short days (SD, 8L:16D, 8h light: 16h dark/dim-light with intensity 1.55:0.002 W/m^2^ respectively) and maintained at a ambient temperature of 22 ± 2°C and monitored using Easy Log USB (Lascar electronics Inc. PA, USA) for 32 weeks. Starting under SD for 4 weeks, birds were either retained in SD (*nonM*) or exposed to long days (LD; 14L:10D). Date of transfer to LD was taken as day 0 of the experiment. Under LD, birds annual life-history states i.e. weeks 1, 5, and 28 were marked as pre-migratory (*preM*, diurnal), migrating (*M*, exhibiting photoinduced night flight behaviour, a simulation of migration in wild conspecifics) and post-migratory (*posM*, diurnal).

### Behavioural and physiological observations

A sub set of the study birds (n=30) were singly housed in activity recording cages (size: 45 cm × 30 cm × 30 cm). A group (n= 6) was maintained as backup to replenish in case of mortality. The rest of the birds (n= 12) were individually housed to record their body weight, surface body temperature, daily food eating pattern, fat score and testicular volume. During nonM and M states, feces were collected for gut microbiota assessment. The Chronobiology Kit software program (Stanford Software Systems, Santa Cruz, CA, USA) was used to collect, plot, and analyse daily activity data. Detailed methodology used for this study were adopted from Gupta et al., 2019, 2020.

### Metabolite assay

Blood samples were drawn from the activity cage housed birds at different states of experiment outlined in Figure 1A. Activity or metabolic alteration due to sampling stress was avoided. Precautions, therein, included-1) reasonable group size (n= 30); 2) a randomized sequence of blood sampling, and 3) a gap of 36 h was followed between two subsequent sampling from the same bird. Starting under SD, blood samples were drawn at 6 time points at 4-hourly intervals, starting lights on i.e. ZT 0, 4, 8, 12, 16 and 20. For this, bird’s wing vein was punctured with a fine sterile needle, and blood droplets (yielding 100-250 μL volume) were gently collected into heparinized microhematocrit capillary tubes. After 4 weeks of SD exposure, 12 birds were retained in SD (*nonM*) and 18 birds were transferred to LD on day 0. Blood samples were drawn from LD birds at *preM, M* and *posM* annual life-history states, respectively. Centrifugation of blood samples, and serum aliquoting was done using methods as in Gupta et al., 2020.

### Quality control (QC) sample and Randomization

An equal volume of all study serum samples (n=144) were pooled to prepare a quality control (QC) sample. Coded study samples were randomized using a web-based tool (www.randomizer.org) and samples were processed in batches of 20 for metabolite extraction and derivatization followed by gas chromatography-mass spectrometry (GC-MS) data acquisition within 24 h of derivatization. To minimize operator biasness, we adopted a double blinding approach during serum sample collection and metabolic profiling study. Two independent researchers carried out sample collection and GC-MS data acquisition.

### Serum Processing and Derivatization

In brief, the serum sample (50 μL) was thawed on ice and freshly prepared isoniazid solution (1 mg/mL, 10 μL) was added as a spike in standard. Ice-cold methanol (800 μL) was added to the sample and vortexed for 30 s. The suspension was centrifuged at 15,000g for 10 min at 4 °C and the supernatant was dried in a SpeedVac at 40 °C. The dried sample was treated with 2% methoxyamine HCl in the pyridine (MOX) reagent at 60 °C for 2 h followed by a silylation step with N,O-bis-(trimethylsilyl)trifluoroacetamide (BSTFA) at 60 °C for 1 h. After derivatization, the sample tubes were centrifuged at 10,000g for 5 min, and the supernatant was transferred into a vial insert kept inside a 2 mL screw-capped glass vial.

### GC-MS Data Acquisition

Using an automated sampler (MPS, Gerstel Germany), the derivatized test serum sample (1 μL) was injected, using the splitless mode, to an RTx-5 column (5% diphenyl, 95% dimethylpolysiloxane; 30 m × 0.25 mm × 0.25 μm; Restek USA) in a GC/TOF-MS (Pegasus 4, Leco, USA). Helium was used as carrier gas at a constant flow rate of 1 mL/min. The inlet temperature was fixed at 300 °C during injection, temperature gradients of 80-150 °C (ramp of 7 °C/ min) and from 150 to 270 °C (ramp of 10 °C/min) with a hold time of 1 min between two ramps. After reaching the final temperature, a hold time of 15 min at the final temperature was maintained. The electron ionization (EI) mode was fixed at 70 eV to scan ions of 35-600 m/z ranges. The maximum scan speed was 20 Hz with a 230 s solvent delay. The ion source temperature was fixed at 240 °C, and data acquisition voltage was 1600. Sample introduction to data acquisition parameters in GC-MS was controlled through ChromaTOF software (Leco, USA), and the total run time was 2340 s per sample.

### Data pre-processing and peak alignment

Raw GC-MS data (.peg) files of all the study samples (n = 144) were pre-processed using ChromaTOF. Alignment of all GC-MS data files was carried out using ‘Statistical Compare’ feature of ChromaTOF. For peak picking, peak width was set at 1 sec and signal/noise (S/N) threshold was 100. For tentative molecular feature identification mainlib (2,12,961 spectra) and replib (30,932 spectra) libraries from NIST (version 11.0) were used with a minimum similarity index of 750. Maximum retention time deference was set at 0.5 sec and for mass spectral match minimum spectral similarity among aligned molecules was set at 600. Unsilylated molecules were removed from the data matrix manually. Aligned peak information was exported to .csv format and molecules absent in more than 50% of samples of at least one class (time point) were excluded from analysis. For metabolite analysis, both uni- and multi-variate analyses of the total area normalized metadata were carried out using MetaboAnalyst 4.0 to identify important deregulated molecules (Xia et al., 2015). Missing values were imputed with half of the minimum value of study population and data matrix was normalized with peak area of the feature representing the internal standard isoniazid. Following which, generalized log transformation and universal scaling method were employed to obtain a near Gaussian distribution of the variables to carry out multivariate analyses (principal component analysis: PCA, and random forest: RF). Univariate analysis in terms of paired t-test (P < 0.05) was carried out using MetaboAnalyst. Features qualifying criteria of t-test (P value < 0.05) and Mean Decrease Accuracy score > 0.002 of a Random Forest (RF) model were selected as important deregulated molecules. Selected important molecular list was taken for hierarchical clustering using Ward linkage and Euclidean cluster method and separately Principal Component Analysis (PCA) was carried out to see the clustering pattern between study groups.

### Confirmation of identity of important molecules

Identity of 47 molecules was confirmed using reported retention indices. Commercial standards of tentatively identified important molecules were oximated and silylated following similar methods used for analysing the samples. Derivatized standards were run following the GS-MS methods used for sample data acquisition. Retention time and fragmentation pattern of both commercial standards and important metabolites were matched to establish the identity of these important molecules.

### Pathway analysis

To identify deregulated molecular pathways, identified important deregulated/all molecules were used as input list in MetaboAnalyst 4.0 (Xia et al., 2015). KEGG pathway 1100 was used for pathway analysis using *Gallus gallus* library.

### Gut microbiota assessment of nonM and M fecal samples

Faecal sample were collected using an indigenous collection kit comprising plastic box with gauze, plastic tray, sterile swabs, and sterile snap top collection tubes. Once the birds defecated, faeces were collected in the tube and quickly stored at 4°C. Notably, storage conditions do not overshadow between-samples differences up to 2 weeks, even at room temperatures (Lauber et al., 2010). Our analyses focused on bacterial composition of faeces rather than abundance. For this, all faeces samples were streaked on individual Luria Bertani Agar (LA, Himedia Inc.) culture plates, prepared following the manufacturer’s protocol and incubated for 40 hours at 37°C. From the culture, quality of DNA was evaluated on 1.0% Agarose *Gel, to observe single band of high-molecular weight DNA*. Fragment of 16S rDNA gene was amplified by 27F and 1492R primers. A single discrete PCR amplicon band of 1500 bp was observed when resolved on Agarose gel. The PCR amplicon was purified to remove contaminants. Forward and reverse DNA sequencing reaction of PCR amplicon was carried out with forward primer and reverse primers using BDT v3.1 Cycle sequencing kit on ABI 3730xl Genetic Analyzer. Consensus sequence of 16S rDNA gene was generated from forward and reverse sequence data using aligner software. The 16S rDNA gene sequence was used to carry out BLAST with the database of NCBI GenBank database. Based on maximum identity score first ten sequences were selected and aligned using multiple alignment software program Clustal W. Distance matrix was generated and the phylogenetic tree was constructed using MEGA 7 (Kumar et al., 2016).

### Statistical analysis

In behavioural study, student’s t-test was used to compare total activity and food intake between different observations. One-way ANOVA followed by post-hoc Newman-Keuls was used for comparing physiological and metabolite concentrations in *nonM, preM, M* and *posM* states at different time points. Significance was selected between observations with a P < 0.05. Data were plotted and statistical analyses performed using Prism GraphPad software (GraphPad ver. 8.0, San Diego, CA).

## Declaration

All authors declare no conflict of interest.

## Acknowledgements

Financial assistance from SERB, New Delhi (letter no. CRG/2019/002542) and UP Centre of excellence (letter no. 78/2022/1984/SATTAR-4-2022-003-70-4099/7/022 S.No.21) is gratefully acknowledged. Permission from Wildlife department Jaipur to work on buntings is gratefully acknowledged. RKN acknowledges the CORE support from ICGEB New Delhi Component.

## Notes

### Competing Interest Statement

The authors have declared no competing interest.

