## Supplementary Figure S1, S2,S3 for "Seasonal plasticity and energy efficiency in migratory buntings is an assemblage of alterations in metabolic and gut microbe adaptations"

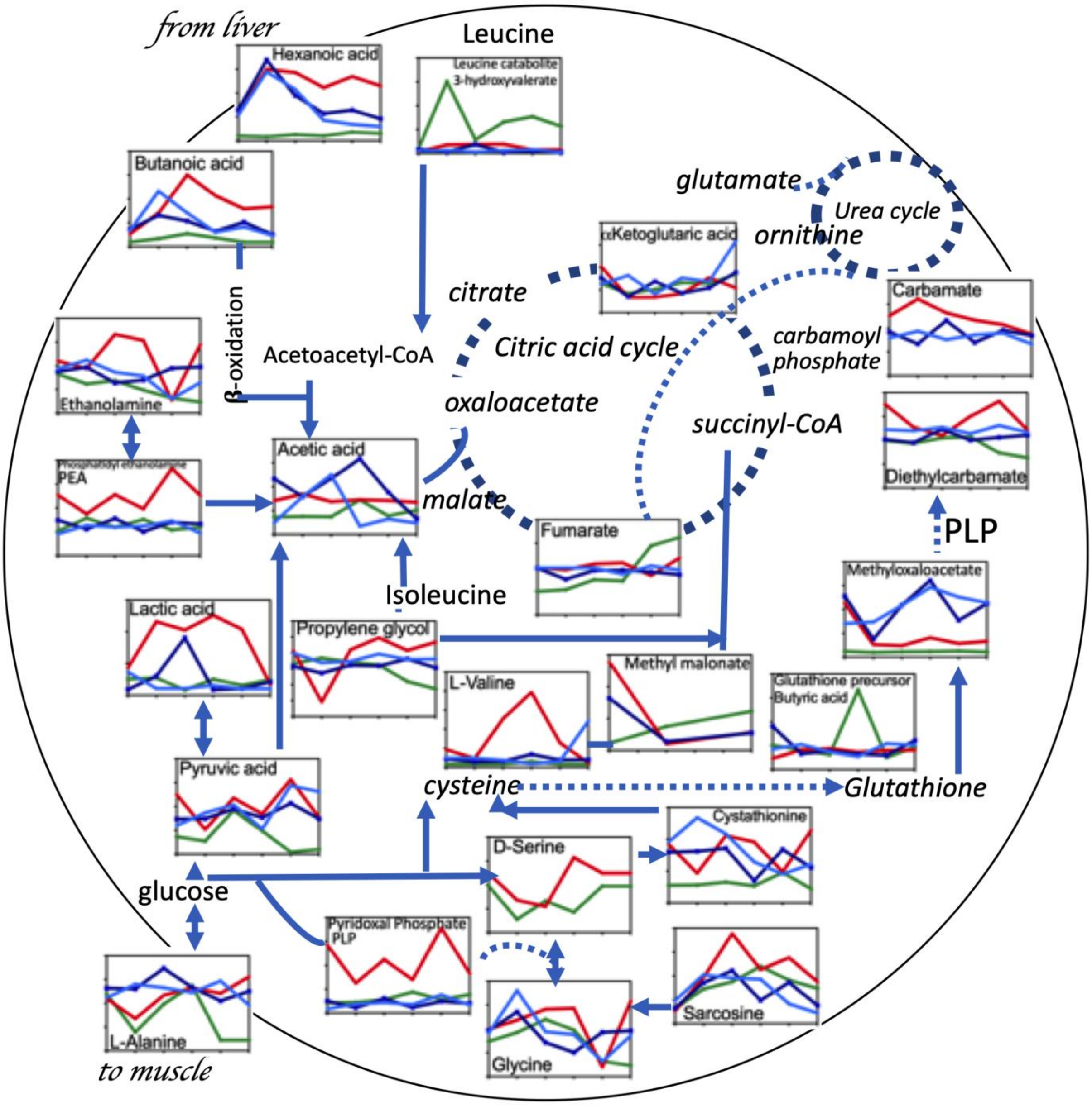


**Supplementary Figure S1.** Daily profile of serum metabolites during nonM (cyan blue, non-migratory), preM (blue, premigratory), M (red, migratory) and posM (green, postmigratory) states. The metabolite graphs are superimposed on metabolite pathway maps of glycolysis, the tricarboxylic acid (TCA) cycle, fatty acid degradation and amino acid metabolism. All P values $*p < 0.05$, $**p < 0.01$ are mentioned in text to retain simplicity of the message in figure.

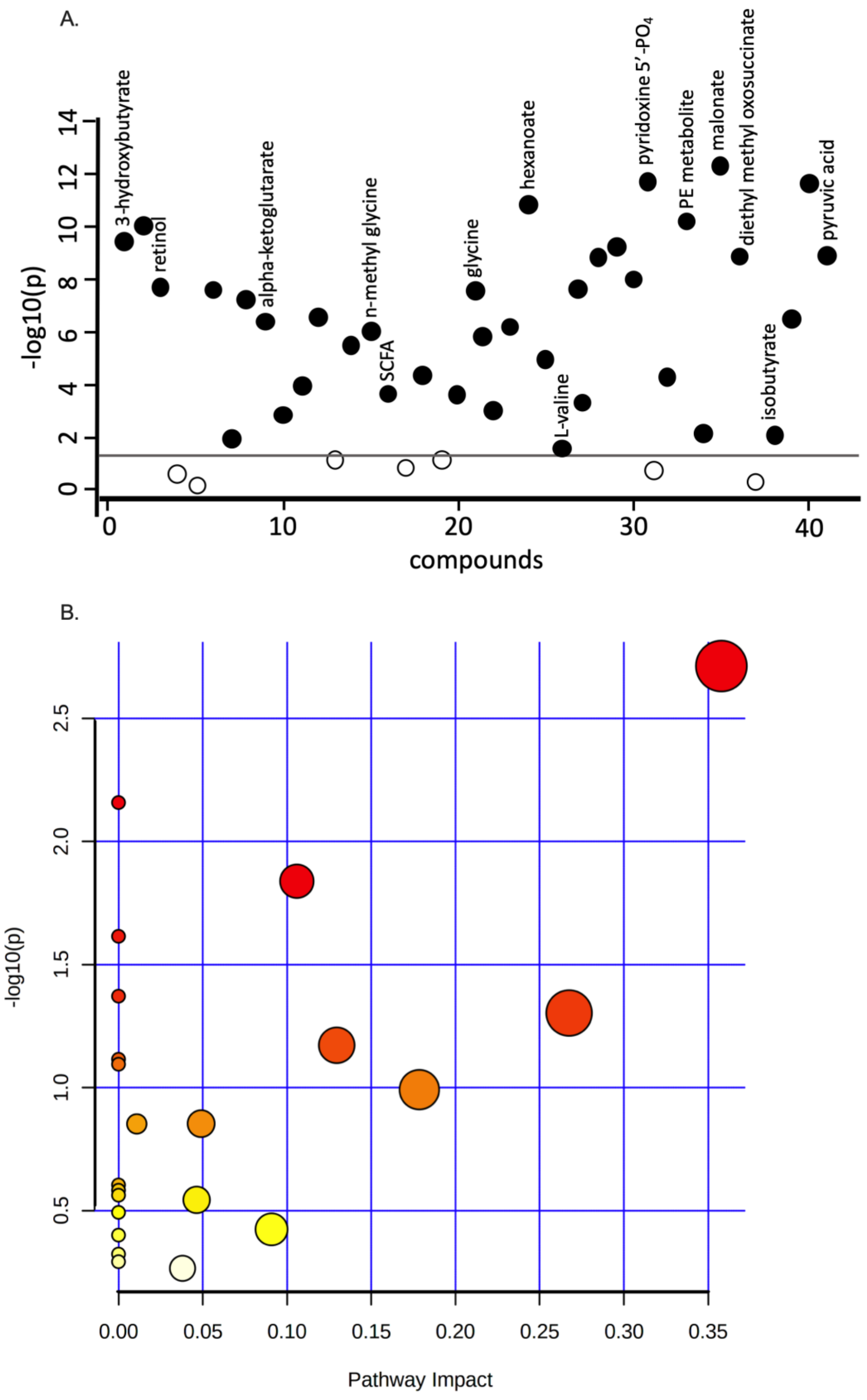


**Supplementary Figure S2.** A. Serum analytes showing significant variation based on fold-change against $-\log_{10}$ (P value). B. Pathway enrichment plot presenting important serum analytes.

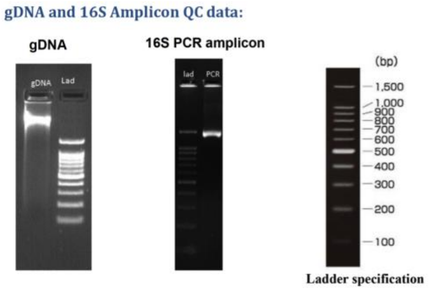


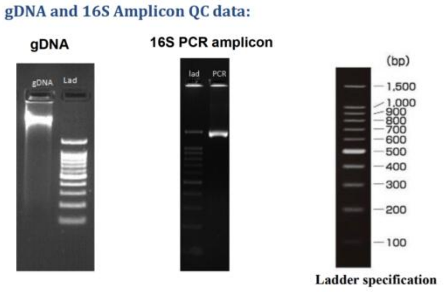


### A. Non-migratory bird gut microbiota

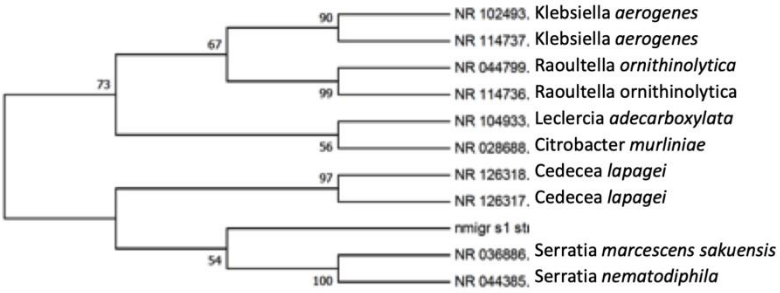


### B. Migratory bird gut microbiota

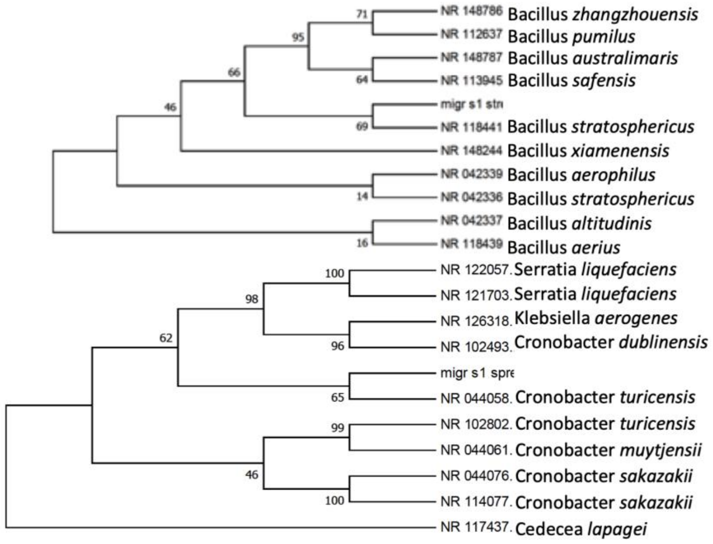


**Supplementary Figure S3.** A. Phylogenetic tree of gut microbiota of non-migratory birds generated by maximum likelihood (ML) analysis of the 16S rDNA sequences. B. Phylogenetic tree of gut microbiota of migratory birds generated by maximum likelihood (ML) analysis of the 16S rDNA sequences.
